# No evidence that portion size influences food consumption in male Sprague Dawley rats

**DOI:** 10.1101/524272

**Authors:** Fabien Naneix, Sophie C. Pinder, Megan Y. Summers, Renee M. Rouleau, Eric Robinson, Kevin P. Myers, James E. McCutcheon

## Abstract

In studies of eating behavior that have been conducted in humans, the tendency to consume more when given larger portions of food, known as the portion size effect (PSE), is one of the most robust and widely replicated findings. Despite this, the mechanisms that underpin it are still unknown. In particular, it is unclear whether the PSE arises from higher-order social and cognitive processes that are unique to humans or, instead, reflects more fundamental processes that drive feeding, such as conditioned food-seeking. Importantly, studies in rodents and other animals have yet to show convincing evidence of a PSE. In this series of studies, we used several methods to test for a PSE in adult male Sprague Dawley rats. Our approaches included using visually identifiable portions of a palatable food; training on a plate cleaning procedure; providing portion sizes of food pellets that were signaled by auditory and visual food-predictive cues; providing food with amorphous shape properties; and providing standard chow diet portions in home cages. In none of these manipulations did larger portions increase food intake. In summary, our data provide no evidence that a PSE is present in male Sprague Dawley rats, and if it is, it is more nuanced, dependent on experimental procedure, and/or smaller in size than it is in humans. In turn, these findings suggest that the widely-replicated PSE in humans may be more likely to reflect higher-order cognitive and social processes than fundamental conditioned behaviors.

**Highlights:**

- Portion size effect (PSE) refers to increased food intake induced by large portions.
- Although widely replicated in human feeding studies, it may not exist animals.
- Presence of a PSE in animals would shed light on mechanisms, which are not known.
- Here, we find no evidence of PSE in male Sprague Dawley rats under a number of experimental conditions.
- This suggests that the human PSE is more likely due to socio-cognitive processes.

## 1. INTRODUCTION

The “portion size effect” (PSE) refers to the increase in energy intake that occurs when an individual is presented with a larger vs. smaller food portions [1]. In humans, this has been widely replicated in both laboratory and naturalistic settings, in both children and adults, using a large variety of foods and experimental protocols [2–5]. Importantly, some have suggested that the increased portion size of food in many societies may, in part, be an important causal factor in the global overeating and obesity crisis [6]. Although the biological and cognitive processes that underpin the PSE are still poorly understood, several potential mechanisms have been suggested. These include: (i) the role of visual cues, including amount of food, food size and unit; (ii) physical properties of food influencing meal structure; (iii) previous experience and learned responses; or (iv) social influence and norms of appropriateness (for review see [4, 5]. In particular, it has been suggested that the PSE could be driven by a learnt tendency to clear one’s plate when eating [7] and/or portion size may signal what a socially ‘appropriate amount’ to eat is [8].

Despite the abundant evidence for a PSE in humans, food portion size as a driver of eating behaviour is rarely considered in animal studies, so it is unknown whether an analogous effect exists in lab animal models. Indeed, a robust PSE has been shown even in people with very different social norms for eating who live an unmodernized lifestyle outside of the Western obesogenic food environment [9].This suggests the possibility that the PSE reflects a basic feature of dietary psychology driven by relatively low level mechanisms, such as conditioning, which could operate similarly in other species. However, most protocols measuring animal feeding behavior tend to use *ad libitum* or an excessive amount of food, relative to daily intake. While this may promote overconsumption, the total amount of food available may exceed the operating range of any analogous PSE. Furthermore, in experiments measuring discrete meals, food is often presented in a way that completely obscures portion size cues (*e.g.* drinking bottle with stainless steel sipper) or using a constant portion size (*e.g.* individual food pellet delivery).

Although the existence of a PSE in laboratory animal models remains unknown, numerous studies have demonstrated that food-related cues, including food itself, can induce food seeking behaviors even in satiated animals and can increase intake [10–14]. Interestingly, some studies have shown that simply increasing the number of sources of palatable food available (e.g. bottles of sucrose solution) is sufficient to increase consumption and weight gain [15]. While these results are not necessarily comparable to a PSE observed in humans (*e.g.* amounts available in a single meal), they strongly suggest that animals, including rodents, are sensitive to external food-related parameters that reflect food abundance and respond to them by increasing their consumption.

Understanding whether non-human species of animals demonstrate a PSE analogous to that seen in humans will help develop better understanding of the drivers of species-specific eating behavior and, in addition, may reveal commonalities with human eating behavior. For example, the influence that portion size has on eating behavior may not be uniquely human and therefore not reliant on higher-order cognition (as a social appropriateness account of the PSE would suggest) and instead a tendency that is observable in other non-human animals. In this series of experiments, we used a range of protocols to investigate the existence of a PSE in rats. These studies were conducted independently across two different laboratories in different countries – the US and the UK – using the Sprague Dawley (SD) strain. Our experiments used: (i) visually identifiable portion size of a palatable liquid food (**Exp 1a**); (ii) initial training on a “plate cleaning” procedure (**Exp 1b**); (iii) portion sizes of palatable pellets signaled by auditory and visual food delivery-associated cues (**Exp 2a**); (iv) amorphous shape properties of food (peanut butter; **Exp 2b**); and (v) standard (chow) diet portions provided in the rats’ natural feeding environment (**Exp 2c**).

## 2. MATERIALS AND METHODS

### 2.1. Experiment 1a

#### 2.1.1. Subjects

Given that our studies were exploratory in nature we based our sample sizes (largest n = 16, smallest n = 8) on being able to detect statistically large effects of PS on food consumption and if any study produced a pattern of results consistent with a PS effect we intended to replicate the finding in a larger sample size. In experiment 1a, male adult SD rats (Charles River laboratories, n = 12) weighing 350-450 g were tested. Rats were housed individually in 20.3 × 40.6 × 26.7 cm plastic tub cages with corncob bedding and maintained at approximately 21°C and 40% humidity, with a 12 h light/dark cycle (lights on at 8:00 AM). All testing occurred in the light phase. Rats had *ad libitum* drinking water and were provisioned with daily chow rations (16 g/day Mazuri Rodent Diet 5663; 3.41 kcal/g, 14% energy from fat, 27% from protein, 59% from carbohydrate) in addition to test meals described below. Procedures of Experiment 1a and 1b were approved by the Bucknell University IACUC and were conducted in accordance with the NIH Guide for Care and Use of Laboratory Animals, 8^th^ edition.

#### 2.1.2. Procedure

First, for 7 days, rats were acclimated to receive a daily chow ration at approximately 6 PM. After this acclimation phase, rats received daily 30-min access to Ensure liquid diet (chocolate Ensure, Abbot Nutrition, Columbus OH) at a pseudo-randomly determined time between 10 AM and 5 PM for 6 consecutive days. Ensure liquid diet was provided in a glass bottle fitted with a rubber stopper and stainless steel sipper tube placed on the individual home-cage lid. Intake was measured by weight. The average of the last four days was considered each rat’s baseline consumption.

Following baseline measurements, rats’ intake was measured in a series of daily tests where portion size was made salient. In these tests, rats were provided with Ensure in a polycarbonate cup (7 cm diameter, 90 ml capacity) affixed to a hanger that could be positioned inside the rat’s home cage. On different days, each rat was provided with either 150% or 200% of its baseline intake. Those two portion sizes were tested twice for each rat in a randomly assigned sequence, counterbalanced across rats for four days. Testing occurred daily at either 10 AM or 4 PM (once each for each portion size) and each rat’s intake was measured by weight of the cup before and after the 30 min test. The average of the two repetitions of each portion was used for analysis. No rat finished the entire amount provided in any of the tests.

### 2.2. Experiment 1b

#### 2.2.1. Subjects

Male adult SD rats (Charles River laboratories, n = 16) weighing 350-450 g were tested. Housing conditions were as described in Experiment 1a.

#### 2.2.2. Procedure

Individual daily energy intake was determined by measuring *ad libitum* daily chow intake during 5 consecutive days (chow was as described in Experiment 1a). For the remainder of the experiment, rats were restricted each day to 90% of that baseline. After 6 days restricted chow rations, the training phase began wherein rats were fed a “snack” portion of chocolate Ensure three times per day for 10 consecutive days. Each rat’s snack portion was 15% of its baseline daily energy intake (mean = 9.2 ± SD 1.23 g of Ensure), and was provided in a polycarbonate cup hung inside the home cage. The three daily snacks were given between 8–10 AM, 12–2 PM, and 4–6 PM. By the end of the second training day, all rats consumed the entire snack portion within 30 minutes of it being provided, with the exception of a single rat who failed to finish the snack portion on three subsequent occasions throughout training. Chow rations constituting the remaining daily energy intake allowance were provided at 6 PM daily.

Following the training phase, rats were tested in a series of 30-min daily intake tests following overnight food restriction. In these tests, rats were given a *Small* (3 times the size of the snack portion, mean = 27.5 ± SD 3.7 g), *Medium* (150% of Small portion; 4.5 times the size of the snack portion, mean = 41.2 ± SD 5.5 g), or *Large* (200% of Small portion; 6 times the size of the snack portion, mean = 54.5 ± SD 7.4 g) portion of Ensure. Rats were given one test meal daily between 12-2PM. The chow rations to be delivered at 6PM were adjusted for kcal consumed in the test meal so that daily kcal remained constant. Each rat was tested twice with each portion size in counterbalanced order, with the average of the two repetitions used for analysis.

### 2.3. Experiment 2a

#### 2.3.1. Subjects

Eight male adult Sprague Dawley rats (Charles River laboratories) weighing between 250-350 g were used for Experiments 2a, 2b and 2c. Rats were housed in pairs in 46.2 × 40.3 × 40.4 cm ventilated cages and maintained at approximately 21 ± 2°C and 40-50% humidity with a 12 h light/dark cycle (lights on at 7:00 AM). All testing occurred in the light phase. Rats had *ad libitum* access to water and food (EURodent Diet 5LF5, TestDiet; 3.40 kcal/g, 9% energy from fat, 26% from protein, 65% from carbohydrate). Procedures were performed in accordance with the Animals (Scientific Procedures) Act 1986 and carried out under Project License 70/869.

#### 2.3.2. Procedure

After one week of acclimation, rats were trained for three days, in behavioral chambers (30.5 × 24.1 × 21.0 cm; Med Associates), to consume 45 mg food pellets (Dustless Precision Pellets, F0021-A, BioServ) during daily 20-30-min magazine training sessions. Chambers were equipped with a house light, a fan, and a pellet dispenser positioned on the center of the right wall. Thirty food pellets were delivered at pseudorandom intervals (mean inter-pellet interval, 40 ± 15 s) into a custom-designed pellet trough (6 × 6.5 × 2 cm; 3D printed using Open Scad 2015.03 and Ultimaker 2+; design available at https://github.com/mccutcheonlab/3dprints).

The effect of portion size on pellet consumption was measured over the next 3 weeks in 30 min daily sessions. Testing occurred between 9-12 AM and although rats were tested in the same order, exact time of testing varied by up to an hour each day. Portion sizes of 10, 15, 20, 30, 50, 70 and 90 pellets were tested using a Latin-square design. At the start of the test session, the house light and fan turned on and there was a 1-min habituation period, followed by the delivery of the pellets (5 pellets per second). Pellet intake and accompanying behavior was observed using a webcam (Microsoft LifeCam HD-3000) positioned on the ceiling of the cage and Open Broadcaster Software (OBS). If rats ceased eating for >2 minutes, they were removed from the cage, the session was terminated, and total pellet consumption was measured.

### 2.4. Experiment 2b

#### 2.4.1. Subjects

The same rats as in Experiment 2a were used for this experiment. Experiment 2a and 2b were separated by 2 days. Housing conditions were identical.

#### 2.4.2. Procedure

First, a habituation day took place during which rats were allowed to freely consume 10 g of peanut butter (Sun Pat Smooth; 48.8 g fat/14.7 g carbohydrates/24.4 g protein per 100 g) placed in a Petri dish (9 cm diameter) during a 30 min session in the behavioral chambers. Testing occurred between 9-12 AM and although rats were tested in the same order, exact time of testing varied by up to an hour each day. Portion size effect was then tested for four days by presenting differently-sized portions of peanut butter (2.5 g, 5 g, 7.5 g and 10 g) using a Latin-square design. At the start of the test session, rats were placed in the chamber, the house light and fan turned on and there was a 1-min habituation period, followed by the placement of a petri dish of peanut butter into the center of the chamber. Food intake was recorded in exactly the same way as in Experiment 2a. At the end of each session the amount of peanut butter left in the Petri dish was measured. The maximum duration for a test session was 30-min.

### 2.5. Experiment 2c

#### 2.5.1. Subjects

The same rats as in Experiment 2a and 2b were used. Housing conditions were identical. Experiment 2b and 2c were separated by 4 days. Rats were food restricted for four days and received daily portion of their standard diet (chow, 15g/rat).

#### 2.5.2. Procedure

The effect of portion size on chow consumption was measured for four days. Four chow portions were tested using a Latin-square design in a series of daily 30 min sessions. Testing occurred between 9-12 AM and although rats were tested in the same order, exact time of testing varied by up to an hour each day. A typical daily meal for the rats was defined as their daily chow portion (15 g, 100%). In other tests they received 150% (22 g), 200% (30 g) and 300% (45 g) of their daily portion. All testing occurred in the rat’s home cage for a 30 min testing session. As the rats were housed in pairs, one of the rats was removed and placed into another empty cage whilst the other rat completed the consumption test.

### 2.6. Data analysis

Data analyses were conducted using GraphPad Prism 7 and Python. All values are expressed as the mean ± SEM. Food intake was analyzed using one-way repeated measures ANOVA or paired two-tailed Student’s t-test when appropriate with portion size as a within-subjects factor. ANOVA tests were followed by Tukey’s *post hoc* tests when appropriate. Measures of effect size (partial eta squared, η_p_^2^) are stated for each comparison. The alpha risk for rejection of the null hypothesis was fixed at 0.05.

All data files are available at Figshare (https://leicester.figshare.com/articles/Portion_size_effect_in_rats/7598942) and custom Python scripts are available on Github (https://github.com/mccutcheonlab/portionsize).

## 3. RESULTS

### 3.1. Experiment 1a: Male SD rats are not sensitive to portion size of palatable liquid diet

Rats (n = 12) received 150% and 200% of their Ensure baseline intake directly in their home cage. All rats ate the palatable diet. However, the portion size of Ensure available did not affect their food intake, as rats consumed a similar amount (**Figure 1a**; 150%: 22.7 ± 1.7 g; 200%: 21.5 ± 2.1 g; paired t-test t_(11)_ = 1.12, p = 0.29, η_p_^2^ = 0.09).

**Figure 1.**
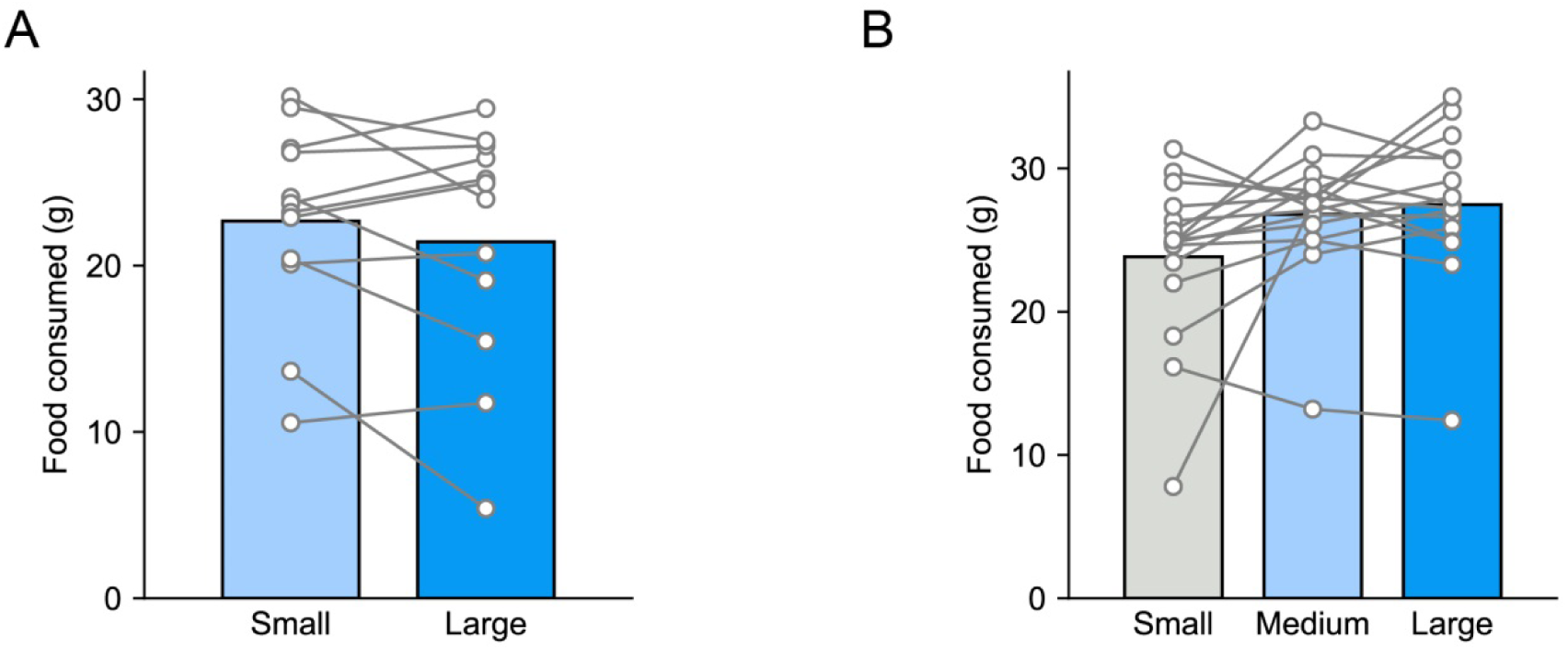
Absence of portion size effect when tested using palatable liquid diet. **(A)** No difference in amount of Ensure intake when rats (n = 12) were given 150% or 200% of their baseline Ensure intake. **(B)** No difference in amount of Ensure intake when rats (n = 16) were given differently sized Ensure portions (3, 4.5 and 6 times the Ensure snack portion). Grey bar (Small) was excluded from analysis because all but three rats consumed the entire portion. Bars show mean intake (in g) and circles are data from individual rats.

### 3.2. Experiment 1b: Male SD rats do not become sensitive to portion size as a result of plate cleaning training

When given access to three times the size of their daily snack Ensure portion (*Small* condition), the majority of rats (13 out of 16, 81%) consumed the entire amount of food available (23.8 ± 1.4 g), thus precluding analysis of this portion. However, no rats consumed the entire amount available when either a *Medium* (4.5 times the snack portion; 26.8 ± 1.1 g) or *Large* (6 times the snack portion; 27.5 ± 1.3 g; **Figure 1b**) portion was provided. Thus, analysis was restricted to the two larger portions and, in this case, food intake between the *Medium* and the *Large* portion size did not differ significantly (paired t-test; t_(15)_ = 0.85, p = 0.41, η_p_^2^ = 0.05).

### 3.3. Experiment 2a: Male SD rats are not sensitive to portion-related visual and auditory food-associated cues

When different numbers of palatable food pellets were delivered in behavioral chambers, rats increased their consumption between 10 pellet and 20 pellet portions but maintained similar consumption levels for the 20-90 pellets portions (**Figure 2**). However, similarly to the situation in Experiment 1b, it was noted that a majority of rats ate the entire 10 and 15 pellet portion, creating a ceiling effect in those tests. Therefore, we confined the analysis to portions from 20 to 90 pellets. Repeated-measures one-way ANOVA revealed no significant effect of pellet portion size on food intake (F_(4,28)_ = 0.21, p = 0.9, η_p_^2^ = 0.03).

**Figure 2.**
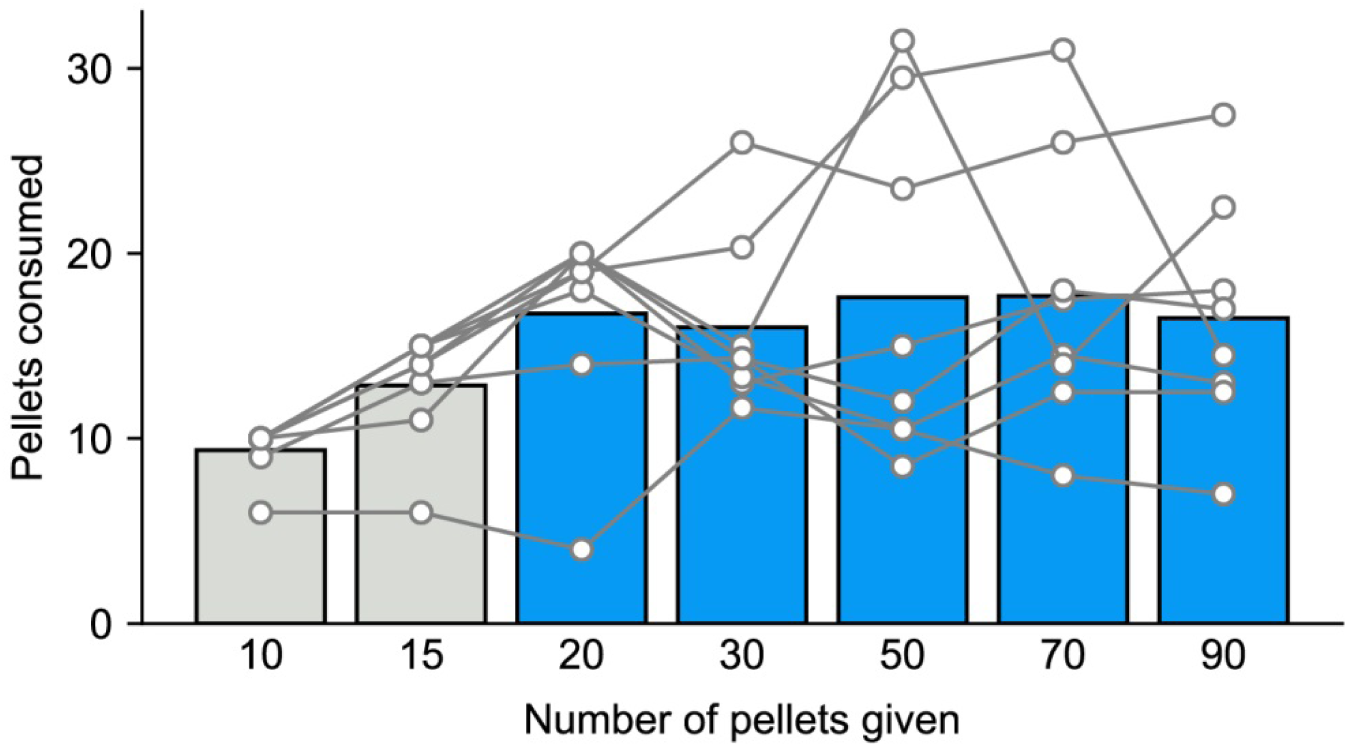
No difference in the number of pellets consumed when rats were given different numbers of pellets (from 10 to 90). Each measurement is the mean of 2-3 replicates for each rat. Grey bars (10 and 15 pellets) were excluded from final analysis because the majority (75%) of rats ate all pellets. Bars show mean pellet intake and circles are data from individual rats.

### 3.4. Experiment 2b: Male SD rats are not sensitive to portion size of an amorphous food

Using the same cohort of rats, we next tested consumption of another type of highly palatable reward, peanut butter, when presented in different-sized portions in a Petri dish directly in the behavioral chambers (**Figure 3**). Rats consumed similar amounts of peanut butter for all the four portion sizes (repeated-measures one-way ANOVA; F_(3,21)_=0.98, p = 0.4). Importantly, in this study, none of rats consumed all of the peanut butter for each portion provided. In addition, we analyzed the time that each rat spent eating in each session (Table 1) but found no significant effect of portion size on this parameter (repeated-measures one-way ANOVA; F_(3,21)_=0.45, p = 0.7, η_p_^2^ = 0.12).

**Table 1.** Time taken spent eating peanut butter when provided with different portions ((i.e. ‘meal’ length; 2.5 to 10 g). Data on total consumption are shown in Figure 3. Values are mean (n=8) with SD in parentheses.

| Peanut butter given (in g) | 2.5 | 5 | 7.5 | 10 |
| --- | --- | --- | --- | --- |
| Time spent eating (in s) | 189.3 (78.2) | 242.1 (170.4) | 232.2 (125.2) | 250.1 (115.1) |

**Figure 3.**
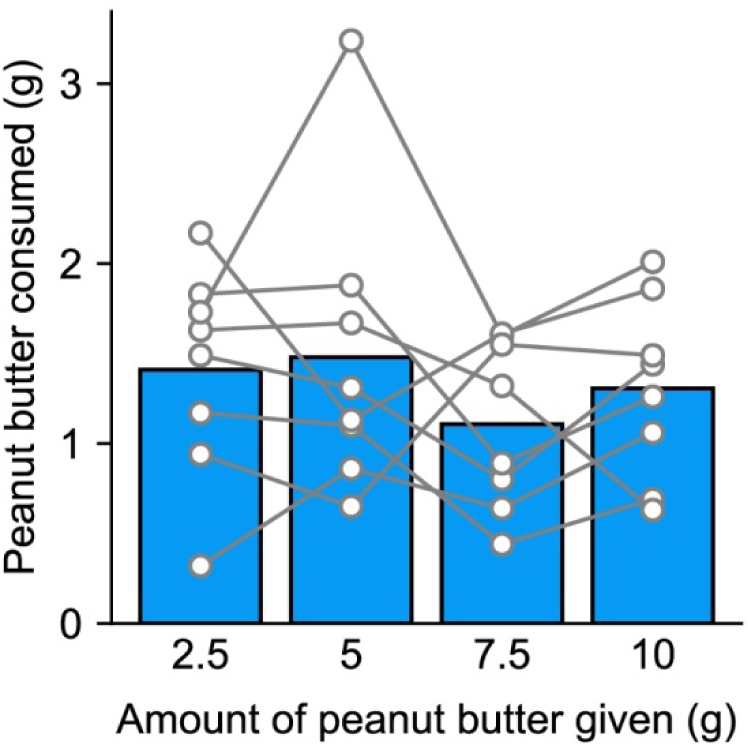
No difference in food intake when rats were given different amounts of peanut butter (from 2.5 to 10 g). Bars show mean intake for all rats (in g) and circles are data from individual rats.

### 3.5. Experiment 2c: Male SD rats are not sensitive to portion size of their standard diet

In a final experiment using the same cohort of rats as tested in Experiment 2a and 2b, we tested the effect of different portions of rats’ standard chow diet provided in the home cage on food intake. Chow consumption was similar between the 15 g portion (6.0 ± 0.5 g) and 45 g portion (6.8 ± 0.6 g; **Figure 4**). Repeated-measures one-way ANOVA confirmed there was no significant effect of chow portion size on food intake (F_(3,21)_ = 0.49, p=0.7, η_p_^2^ = 0.07).

**Figure 4.**
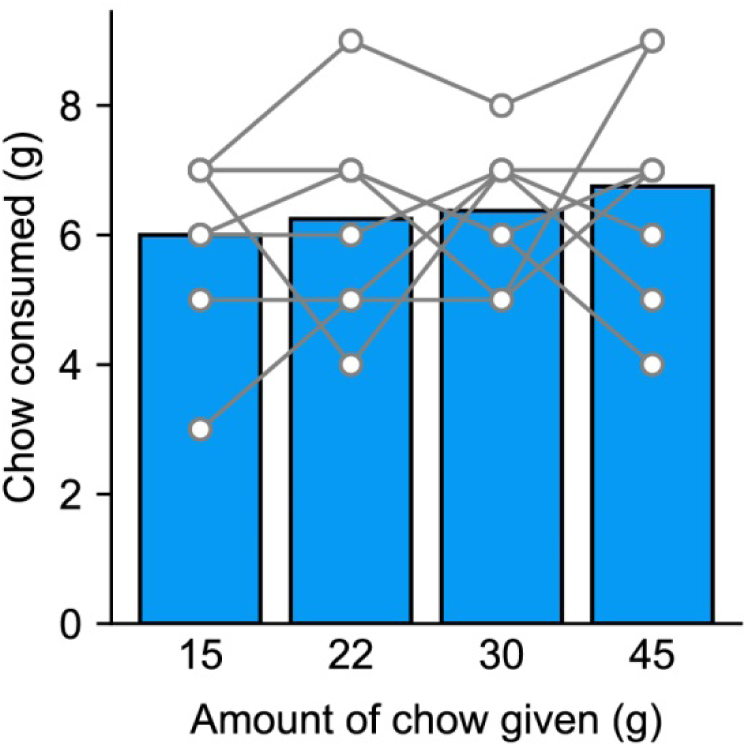
No difference in the amount of chow consumed in home cages when rats were given different amounts (from 15 to 45 g). Bars show mean intake for all rats (in g) and circles are data from individual rats.

## 4. DISCUSSION

The current study investigated if a PSE similar to the one commonly seen in humans was observable in a rodent model. By using five protocols investigating different parameters that could potentially underpin the PSE (quantity of reward, plate cleaning training, visual and auditory cues, shape of the food, familiar food), we found no convincing evidence that male Sprague Dawley rats are sensitive to the size of food portion.

Processes underlying PSE in humans remain poorly understood. Several theories suggest that PSE may be related to the response to different food-related cues including sensory properties of the food itself, dishware size, or the size of the food unit (for review see [4, 5]). Moreover, numerous studies have demonstrated in humans and rodents alike that food-predictive cues in the environment are able to elicit preparatory responses that promote food-seeking [10–12, 16]. Accordingly, food-cue reactivity could promote PSE by inducing increased reactivity to increased stimulus presentations (e.g. sensory cues associated with food delivery). In the present study, all included visually perceptible cues associated with food delivery and that scaled with the amount of food available. Our results, however, demonstrate that food intake of rats is not sensitive to the absolute amount of food provided despite visual cues being available. Moreover, the absence of this effect is not dependent on a specific type of food as we observed similar results with liquid diet (Ensure solution; Experiment 1), small pellets (Experiment 2a), amorphous food (peanut butter; Experiment 2b), or standard rodent chow (Experiment 2c).

As visual perception does not represent the main sensory modality of rodents, Experiment 2a was designed to add auditory cues to the visual perception of the food amount. In this test, food pellets were not present at the beginning of the session and were delivered after 1-min habituation to the behavioral chamber. The sound of pellet dispenser and of the pellets dropping into the trough can be used to identify the portion size (number of pellets delivered). However, as for the other tests, we did not observe difference in food intake, suggesting that neither visual or auditory cues, nor a combination, are sufficient to induce a PSE in our rats. These conclusions notwithstanding, it is notable that in our experiments we do not know whether rats were able to effectively discriminate between the portion sizes we presented. Thus, although we attempted to make the differences as salient as possible by using visual, or auditory cues, or a combination of both, it is possible that lack of portion size effect resulted from an inability to discriminate these cues.

A study in human participants demonstrated that a PSE can be observed even without visual perception of food during eating [17] suggesting that on-going visual cues whilst eating may not be necessary to drive the PSE and that pre-meal awareness of the amount of food available may be of most importance [1]. Furthermore several results suggest that the physical properties of the food itself affect meal macro-and microstructure and may participate in the PSE [18]. Specifically, it has been suggested that portions of food that are amorphous in shape, such as macaroni cheese, are difficult to judge in size and this could support increased portion size consumed[19]. In the present study, Experiment 1 and Experiment 2b used amorphous shape food (Ensure solution and peanut butter). However, we did not observe significant PSE in animals when these foods were presented in excessive amount. Moreover, we also observed that the different amount of peanut butter available (Experiment 2b) does not affect the mean meal duration suggesting no effect of the portion size on meal macrostructure.

It is also the case that in each of the procedures we conducted, the olfactory stimulus arising from the food portion, which is likely a more ecologically significant cue for rats, would presumably scale with portion size, although we were not able to precisely control or quantify that variable. For rats, the intensity of the olfactory cues when a test meal is delivered and/or the continued presence of olfactory stimulus as a meal progresses could be analogous to the role of visual perception in humans. Yet our results do not support the idea that the magnitude of olfactory stimulation arising from the food is, on its own, able to induce the PSE in SD rats.

Dishware emptiness and previous training to habitual “plate clearing” is another possible mechanism proposed for the PSE[7]. It is possible that plate cleaning tendencies may come from socialization and educational instruction [20, 21] that could ultimately result in greater external situational cues (such as portion size) instead of internal signals such as hunger and satiety [4, 22, 23]. In Experiment 1b, rats received initial training to clear a ‘dish’ filled with a snack portion of Ensure solution before being tested with an increased amount of the food presented on the same ‘dish’. The training was intended to make the sensory properties emanating from the food present in the dish as a salient, appetitive cue that could potentially drive subsequent food-seeking. However, neither the presentation of medium (4.5 times the snack portion) nor large (6 times the snack portion) induced an increase in food intake. Although these data suggest that previous training does not induce increase food intake with larger portion, we cannot totally exclude this possibility as the present study involved a limited amount of initial training.

Human and rat eating habits vary considerably. When eating, humans have the cognitive capability to be aware of how socially appropriate their eating behavior is and it has been proposed that higher-order cognitions like this may explain the PSE [4, 8]. Rats are grazers and will on average consume ten meals a day of 2-3 g [24], whereas humans, typically consume 3 meals [25]. This difference may have a major impact on the quantity of food consumed in a single meal across species. It may be the case that if the PSE is observed among non-human animals, it would only be observed among those that tend to eat distinct and less frequent ‘meals’. As such, maximizing food intake (i.e. being responsive to the total amount of food available) under these conditions may be adaptive. The difference in eating frequency between rodents and humans could in theory be circumvented by altering the feeding habits of rats using time-restricted or portion-restricted feeding. In the present study, three of the five tests were performed in food-restricted animals that had access to food only during one or two meals per day. In these studies, rats’ motivation to consume at each opportunity would also be enhanced potentially making the situation more similar to that seen in humans at mealtime. However, even under food-restricted conditions we did not observe a PSE. One observation that we have not been able to exclude is that circadian rhythm could have an effect on PSE. In our study, all tests were conducted during the light phase whereas the majority of rodent food consumption takes place during the dark phase [24]. Thus, the existence of PSE in rodents during their active phase remains to be investigated.

A limitation of our studies was that our sample sizes tended to be small because of their exploratory nature and therefore results of individual experiments should be interpreted accordingly. However, the pattern of results across all five experiments was consistent and, irrespective of the statistical power of inferential analyses, there was no obvious trend across studies suggesting that portion size affected amount of food eaten and that an effect would be detectable with much larger sample sizes. Both in Experiment 1b and 2a we noticed an increase in consumption between the smallest portion (Small portion, Exp. 1b; 10 and 15 pellets, Exp. 2a) and the other food portion. However, this increase must be interpreted carefully as most rats consumed this portion in its entirety thus confounding a potential PSE with a ceiling effect. The only possible exception to this was in Study 2c, in which a 200% increase in portion size of rats’ daily diet was associated with a statistically non-significant 13% increase in consumption across rats, although we note that only 4 of the 8 rats tested increased their food consumption in response to the 200% increase in portion size. This non-significant finding may therefore reflect random fluctuation in food consumption, but may warrant further attention.

Finally, these studies were conducted in two different laboratories on two different continents and for consistency we used male SD rats in both settings. It is possible, that the lack of effect in our hands is either strain or sex-specific and that replicating these experiments in a different strain or in female rats would uncover evidence of a PSE. However, it should be noted that in humans PSE has been widely reported in both sexes and across many varied populations of subjects [9, 17, 19, 26, 27].

To conclude, the aim of this study was to establish whether PSE exists in laboratory rats. In our five consumption tests we failed to find evidence for a PSE in male SD rats suggesting that either portion size does not affect their feeding behavior or that it can only be observed in certain, nuanced conditions. This issue requires further experiments to in which multiple strains of male and female rats, mice and other mammals are compared. An intriguing possibility is that PSE is a process specific to humans. The reason for this is not clear but may reinforce the importance of top-down cognitive processes in driving food intake in humans, a factor that is likely key to understanding the current health crisis of overeating and obesity.

## Author’s contribution

FN, SP, MS, RR, KM and JM performed experiments and analyzed data; FN, SP, ER, KM and JM drafted and edited the final manuscript.

## 5. AKNOWLEDGMENTS

The authors acknowledge the help and support from the staff of the Division of Biomedical Services, Preclinical Research Facility, University of Leicester, and the animal caretaking staff at Bucknell University for technical support and the care of experimental animals.

## 6. FUNDING

This work was supported by the Biotechnology and Biological Sciences Research Council [grant # BB/M007391/1 to JEM]; and the European Commission [grant # GA 631404 to JEM]. ER’s salary is supported by the Medical Research Council (grant MR/N000218/1) and has previously received funding from Unilever and the American Beverage Association.

